# DeepRank-GNN-esm: A Graph Neural Network for Scoring Protein-Protein Models using Protein Language Model

**DOI:** 10.1101/2023.06.22.546080

**Authors:** X. Xu, A. M. J. J. Bonvin

## Abstract

**Motivation:** Protein-Protein interactions (PPIs) play critical roles in numerous cellular processes. By modelling the three-dimensional structures of the correspond protein complexes valuable insights can be obtained, providing, for example, starting points for drug and protein design. One challenge in the modelling process is however the identification of near-native models from the large pool of generated models. To this end we previously developed DeepRank-GNN, a graph neural network that integrates structural and sequence information to enable effective pattern learning at PPI interfaces. Its main features are related to the Position Specific Scoring Matrices (PSSM), which are computationally expensive to generate and significantly limit the algorithm’s usability.

**Results:** We introduce here DeepRank-GNN-esm that includes as additional features protein language model embeddings from the EMS-2 model. We show that the ESM-2 embeddings can actually replace the PSSM features at no cost in-, or even better performance on two PPI-related tasks: scoring docking poses and detecting crystal artifacts. This new DeepRank version bypasses thus the need of generating PSSM, greatly improving the usability of the software and opening new application opportunities for systems for which PSSM profiles cannot be obtained or are irrelevant (e.g. antibody-antigen complexes).

**Availability and implementation:** DeepRank-GNN-esm is freely available from https://github.com/DeepRank/DeepRank-GNN-esm

## 1. Introduction

Protein-protein interactions (PPIs) are essential for many biological processes. Obtaining three-dimensional (3D) structural information on the corresponding assemblies is key to uncover their functions, and misfunctions in case of disease. Experimentally, this is typically done by cryo-electron microscopy or tomography, X-ray crystallography or, to a less extent, Nuclear Magnetic Resonance spectroscopy. This is, however, often a labour-intensive and costly process. As a result, computational approaches, often supplemented by a limited amount of data, have emerged as valuable alternatives. This has been done classically by docking and/or integrative modelling approaches (Koukos and Bonvin, 2020; van Noort *et al*., 2021; Braberg *et al*., 2022). Artificial Intelligence (AI) has also entered the field of complex prediction, in particular with various applications of AlphFold2 (Jumper *et al*., 2021) and Alpha Fold-multimer (Evans *et al*., 2022) to the prediction of protein-protein (Zhu *et al*., 2022; Gao *et al*., 2022) and protein-peptide (Johansson-Åkhe and Wallner, 2022; Chang and Perez, 2023) complexes.

One of the challenges in computational approaches is accurately identifying near-native PPI conformations from the large pool of generated models, which is often referred to as “scoring”. Multiple in-silico approaches have been proposed to this end (Xue *et al*., 2015; Casadio *et al*., 2022; Geng *et al*., 2019), including physics-based scoring function implemented e.g. in HADDOCK (Dominguez et al., 2003), knowledge-based statistical potentials such as GOAP (Zhou and Skolnick, 2011), classical machine learning methods such as iScore (Geng et al., 2020), meta predictors (Jung *et al*., 2023) and, in recent years, deep learning approaches such as DOVE (Wang *et al*., 2020), DeepRank (Renaud *et al*., 2021), GNN-DOVE (Wang *et al*., 2021) and a Graph Neural Network (GNN) version of DeepRank, DeepRank-GNN (Réau *et al*., 2023), which was shown to have the best performance at the time of publication. It was recently applied to the classification of physiological versus non-physiological interfaces in homomeric complexes, showing the best performance of all single predictions (Schweke *et al*., 2023).

DeepRank-GNN converts three dimensional structures of PPI complexes into residue-level graphs and uses the resulting graph interaction network for making predictions. It was trained on two tasks: classification of crystallographic interfaces and scoring of PPI complexes. In both cases, the Position-specific Scoring Matrices (PSSMs) were found to be the main features driving the predictions. PSSMs provide valuable information about the evolutionary conservation profiles of residues at interfaces of PPIs, helping to identify functionally important residues. However, computing PSSMs is computationally expensive, particularly when aiming at larger alignment depths that enhance the results’ reliability. This requirement, coupled with the need for a non-redundant protein sequence database, can pose challenges for users of the software and impact its overall usability. Despite the availability of pre-computed PSSMs at external databases such as 3DCONS (Sanchez-Garcia *et al*., 2017) and Conserved Domain Database (Lu *et al*., 2020), they often have limitations in their content. Therefore, there is a need to find more flexible and fast way to compute features to enhance the usability of DeepRank-GNN.

The scaling of large language models (LLMs) to incorporate billions or even trillions of training parameters has unlocked unprecedented capabilities, enabling advanced reasoning and the generation of lifelike images and text (Wei *et al*., 2022). This transformative progress in natural language processing has not only revolutionized the field but has also paved the way for the development of protein language models. Evolutionary Scale Modeling-2 (ESM-2) is a state-of-the-art transformer architecture trained on over 65 million unique protein sequences to predict the identity of randomly masked amino acids (Lin et al., 2023). By leveraging a massive-scale training approach that involves solving missing puzzles with over 15 billion parameters, ESM-2 is able to effectively internalize complex sequence patterns across evolution and generate high-quality embeddings that are rich in both evolutionary and functional insights. ESM-2 embeddings, which are very fast to compute, are therefore valuable for various protein-related tasks, such as structure prediction, design, and functional annotation (Lin *et al*., 2023).

To bypass the lengthy computation of PSSMs, we present here DeepRank-GNN-esm, a new version of our DeepRank-GNN algorithm that incorporates ESM-2 embeddings. The process of generating ESM-2 embeddings for a protein sequence is significantly more efficient in terms of computational resources and time investment as it does not rely on multiple sequence alignments (MSAs). We show that those can substitute the PSSM features at no loss in performance. By integrating ESM-2 embeddings into DeepRank-GNN, we achieve a significant acceleration in PPI scoring tasks, resulting in improved usability without compromising the performance. We also show that combining PSSMs and ESM-2 embedding does lead to an improvement in overall scoring performance indicating, that they do contain complementary information. Our algorithm is freely available as a Python package (https://github.com/DeepRank/DeepRank-GNN-esm), featuring two re-trained models, making it easily accessible for researchers in the field of structural biology and bioinformatics.

## 2. Materials and methods

### 2.1 Datasets

#### 2.1.1 BM5 and CAPRI datasets

We used the same training and test dataset as previously used for DeepRank and DeepRank-GNN (https://doi.org/10.15785/SBGRID/843). It consists of docked models of 143 complexes from the Docking benchmark dataset version 5 (BM5) (Vreven et al., 2015), excluding antibody-antigen complexes and complexes with more than two chains. To address potential variations in model performance arising from the selection of data subsets for training, as noted by (Réau *et al*., 2023), we randomly divided the 128 complexes and their associated models into training (80%) and evaluation (20%) datasets by complex for cross-validation while reserving 15 complexes as the independent test set. The final models were obtained by training on the full training dataset as aligned with the training methodology used in the final model of DeepRank-GNN. Additionally, as an independent test set we took 13 complexes from the CAPRI score set (Lensink and Wodak, 2014) as described in DeepRank and DeepRank-GNN, details of which can be found in Supplementary Table S1, to further validate our algorithm’s performance.

#### 2.1.2 MANY/DC benchmark

We explored the application of our proposed models on the task of detecting crystal artifacts from true biological interfaces. Classification of biological or crystallographic PPIs is challenging. In a recent community-wide investigation on assigning protein complexes to correct oligomeric state, DeepRank-GNN achieved the highest AUC among 252 scoring functions (Schweke et al., 2023). To further investigate the efficacy of ESM features, we trained two binary classification models on the MANY dataset (Baskaran *et al*., 2014) and assessed their performances on the DC dataset (Duarte *et al*., 2012), following the methodology employed by DeepRank and DeepRank-GNN. The MANY/DC benchmark is accessible from https://doi.org/10.15785/SBGRID/843.

### 2.2 Graph generation

To construct the protein graphs, we selected protein residues located within 8.5 Å of the interface of the complex as graph nodes. Interface edges and internal edges of the graphs were defined as previously described (Réau *et al*., 2023). Node and edge features were computed and stored in HDF5 format for efficient processing. We calculated ESM-2 embeddings for each protein sequence with model and scripts provided by Lin *et al*. and assigned the embedding for each residue to the corresponding graph node. Details of the ESM-2 embeddings computation can be found in the Supplementary Method 1 section.

### 2.3 Model training

We adapted the model architecture of DeepRank-GNN (Réau et al., 2023) and trained three algorithms (DeepRank-GNN-esm-pssm, DeepRank-GNN-esm and DeepRank-GNN-no-pssm) to predict the fraction of native contacts (*f*_*nat*_) for PPI conformations. The various features used in each model are listed in Table 1. During model training, we used a mean squared error loss function, optimized the models using the Adam algorithm with a batch size of 128 and a learning rate of 0.001. We trained all models for 20 epochs. Supplementary Figures S1-S3 show the training and validation loss curves, along with the corresponding AUC values, for both the final models trained on the complete dataset and the models generated during cross-validation.

**Table1.** Model features and the number of total trainable parameters.

| Features<sup>b</sup> | Dimension<sup>c</sup> | Model<sup>a</sup> |  |  |  |
| --- | --- | --- | --- | --- | --- |
|  |  | 1 | 2 | 3 | 4 |
| Type | 20 | ✓ | ✓ | ✓ | ✓ |
| Polarity | 4 | ✓ | ✓ | ✓ | ✓ |
| BSA | 1 | ✓ | ✓ | ✓ | ✓ |
| Charge | 1 | ✓ | ✓ | ✓ | ✓ |
| Cons | 1 | ✓ |  |  | ✓ |
| PSSM_IC | 1 | ✓ |  |  | ✓ |
| PSSM | 20 | ✓ |  |  | ✓ |
| Embedding | 1280 | ✓ | ✓ |  |  |
| Distance | 1 | ✓ | ✓ | ✓ | ✓ |
| Total trainable parameters |  | 52169 | 51465 | 10505 | 11209 |

a) Model 1: DeepRank-GNN-esm-pssm, Model 2: DeepRank-GNN-esm, Model 3: DeepRank-GNN, and Model 4: DeepRank-GNN-no-pssm.
b) Type represents the amino acid type, BSA the buried surface area of the complex, Cons the Conservation score (from PSSM), PSSM IC the Information content (from PSSM), PSSM the Position-specific scoring matrix, distance the normalized distance between graph nodes.
c) Shape represents the dimension of each feature

## 3. Results

### 3.1 Combining PSSMs and ESM embedding improves the scoring performance

#### 3.1.1 Performance of 10-fold cross-validation on the BM5 evaluation set

We trained DeepRank-GNN-esm-pssm, which combines both PSSM and ESM-2 embeddings as features, using 10-fold cross-validation on the BM5 training set. The average AUC obtained across all folds on their evaluation set is 0.639±0.054 (see Supplementary Figure S1). The model performance of the final DeepRank-GNN-esm-pssm model trained on the full training set is evaluated on the independent validation set using AUC and six machine learning metrics: Precision, MCC (Matthews’s correlation coefficient), F1, Recall, R2, and Pearson correlation coefficient (see Supplementary Method 2 for their definition). DeepRank-GNN-esm-pssm model has the highest AUC (0.95) during training (Supplementary Figure S4). Since the dataset is highly imbalanced, we focus our discussion on the F1 score and MCC. The training results show that the DeepRank-GNN-esm-pssm model, which combines PSSMs and ESM embeddings, outperformed the original DeepRank-GNN model with a 7.78% increase in F1 and an 8.40% increase in MCC (Table 2). Moreover, the DeepRank-GNN-esm-pssm model shows a stronger correlation between predicted fnat values and the ground truth than the original model, as indicated by a higher R^2^ value (0.916 vs 0.826). This enhanced correlation is clearly visible in the scatter plot in Supplementary Figure 5.

**Table2.** Comparison of model performances on the BM5 evaluation set.

| Model<sup>a</sup> | Metrics |  |  |  |  |  |
| --- | --- | --- | --- | --- | --- | --- |
|  | Precision | MCC | Recall | F1 | R<sup>2</sup> | Pearson_r |
| 1 | **0.882** | **0.877** | **0.891** | **0.886** | **0.916** | **0.957** |
| 2 | 0.87 | 0.866 | 0.881 | 0.876 | 0.921 | 0.96 |
| 3 | 0.824 | 0.809 | 0.82 | 0.822 | 0.826 | 0.909 |
| 4 | 0.708 | 0.723 | 0.782 | 0.743 | 0.683 | 0.842 |

a) Model 1: DeepRank-GNN-esm-pssm, Model 2: DeepRank-GNN-esm, Model 3: DeepRank-GNN, and Model 4: DeepRank-GNN-no-pssm.

The enhanced performance on the validation set can be attributed to two factors. Firstly, the DeepRank-GNN-esm-pssm model uses the ESM embeddings, which provide both more features (1328) and additional information. The larger feature size results in an increased number of learnable parameters, enhancing the model’s ability to generalize patterns from the input data (Table 1).

#### 3.1.2 Performance on two independent test sets

We first evaluated all 11 DeepRank-GNN-esm-pssm models on the independent BM5 test set by computing the AUC using a cutoff of 0.3 to distinguish between good and bad models (DeepRank-GNN gives a continuous scale output between 0 and 1). To create a smoother curve, the true positive rate (TPR) values are estimated at particular false positive rate (FPR) values using interpolation. The results, depicted in Figure 1, demonstrate that the final model (foldall) achieves the highest AUC value (0.952) compared to all models obtained in cross-validation. Interestingly, eight models trained on subsets of the data outperforms the original DeepRank-GNN model. The variation in performance between folds also highlights the dataset dependency of the algorithm. To be aligned with DeepRank-GNN, we selected the final model trained on the full dataset for subsequent analysis.

**Figure 1:**
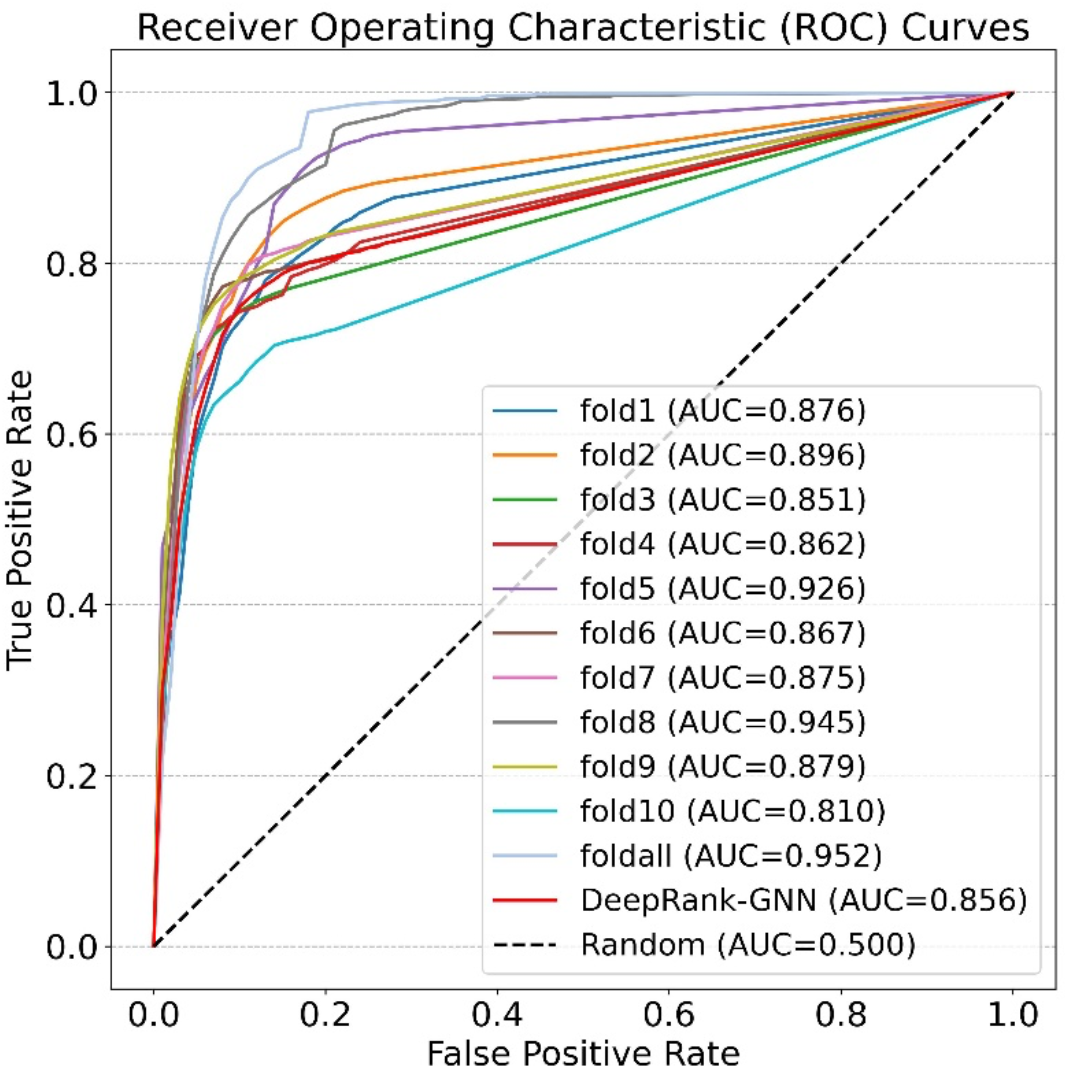
Receiver operating characteristic curves on the BM5 test set. (A true positive is defined as a model with *f*_*nat*_ >0. 3)

As second test set, to ensure a fair comparison of the model’s performance outside of BM5, we compared it with six other scoring methods on the CAPRI score set, including DeepRank (Renaud *et al*., 2021), DeepRank-GNN (Réau *et al*., 2023), GNN_DOVE (Wang *et al*., 2021), HADDOCK (Dominguez *et al*., 2003), iScore (Geng *et al*., 2020) and GOAP (Zhou and Skolnick, 2011). As metric for the comparison, we use the per-complex success rate, which is calculated by counting the number of docking cases in which at least one near-native model is found among the top-k ranking models, divided by the total number of cases. Table 3 demonstrates the superior performance of DeepRank-GNN-esm-pssm over DeepRank-GNN, particularly in relation to the top 5 ranks. DeepRank-GNN-esm-pssm achieves a 46.2% success rate in correctly identifying the structures, surpassing DeepRank-GNN (38.5%). While DeepRank-GNN shows the same performance in scoring near-native conformations when considering a higher number of models, we consider the performance at earlier ranks to be more crucial for PPI scoring tasks. Additionally, DeepRank-GNN-esm-pssm achieves the highest AUC among all the methods

**Table3.** Comparison of model performances on the CAPRI Score set.

| Algorithm<sup>a</sup> | AUC | Success rates (%) |  |  |  |
| --- | --- | --- | --- | --- | --- |
|  |  | Top1 | Top5 | Top20 | Top50 |
| Model1 | **0.81** | 15.4 | 46.2 | 61.5 | 61.5 |
| Model2 | **0.81** | 15.4 | 38.5 | 61.5 | **69.2** |
| Model3 | 0.71 | 7.7 | 38.5 | 61.5 | 61.5 |
| Model4 | 0.69 | 23.1 | 30.8 | 53.8 | 53.8 |
| DeepRank | 0.59 | 15.4 | 15.4 | 53.8 | 61.5 |
| HADDOCK | 0.55 | 23.1 | 23.1 | 46.2 | 61.5 |
| iScore | 0.64 | **38.5** | 46.2 | 53.8 | **69.2** |
| GOAP | 0.42 | 0 | 23.1 | 46.2 | 53.8 |
| GNN-Dove | 0.54 | 15.4 | **53.8** | **69.2** | **69.2** |

a) Model 1: DeepRank-GNN-esm-pssm, Model 2: DeepRank-GNN-esm, Model 3: DeepRank-GNN, and Model 4: DeepRank-GNN-no-pssm.

These results demonstrate the improved scoring performance achieved by combining PSSMs and ESM-2 embeddings in the DeepRank-GNN architecture.

### 3.2 ESM embeddings can substitute PSSM features without any performance loss

#### 3.2.1 Performance of 10-fold cross-validation on the BM5 evaluation set

To further explore the feasibility of substituting the PSSM features with ESM embeddings, we compared the performance of a modified version of our model, termed DeepRank-GNN-esm, which excludes all PSSM-related features (PSSM, PSSM-IC, and Cons). Despite not reaching the same performance level as the DeepRank-GNN-esm-pssm model, the DeepRank-GNN-esm model demonstrated superiority over the original DeepRank-GNN model, with a 7.43% increase in recall and a 7.04% increase in MCC (Table 2). These findings indicate that PSSM features can be substituted by ESM embeddings as an effective alternative which does not require the more costly computation of the PSSM.

Both PSSM and ESM embeddings have the inherent capacity to capture evolutionary conservation, enabling the detection of functionally important residues. ESM embeddings offer a substantial increase in information compared to PSSMs, both in terms of the database involved in the computation process and the dimensionality of the features. This is attributed to the fact that PSSM features, with 20 features per residue, are derived from sequence alignment with the non-redundant (NR) database, whereas ESM embeddings, with 1280 features per residue, are obtained from training on a significantly larger volume of sequences (∼138 million sequences). This distinction potentially explains the effectiveness of substituting PSSM feature with the embeddings. In contrast, the DeepRank-GNN-no-pssm model, which excludes both PSSM and ESM embeddings, has high training loss even after 20 epochs (Supplementary Figure S4), which indicates its limited ability to effectively learn from the input data. This model also shows the lowest performance across all metrics.

#### 3.2.2 Performance on the BM5 and CAPRI test sets

Our comparison of model performance on the BM5 test set demonstrates that the DeepRank-GNN-esm model outperforms the original DeepRank-GNN model in terms of overall AUC. Furthermore, looking at the per-complex AUC for the four models (Figure 2) reveals that the DeepRank-GNN-esm model achieves the highest AUC of 0.799, surpassing the original DeepRank-GNN model (0.649) by over 20%. The DeepRank-GNN-no-pssm model, which excludes both PSSM and ESM-2 embeddings, exhibits the lowest AUC value (0.589). It is however important to acknowledge that all four models display relatively high standard deviations, suggesting potential challenges in generalization across all target classes.

**Figure2.**
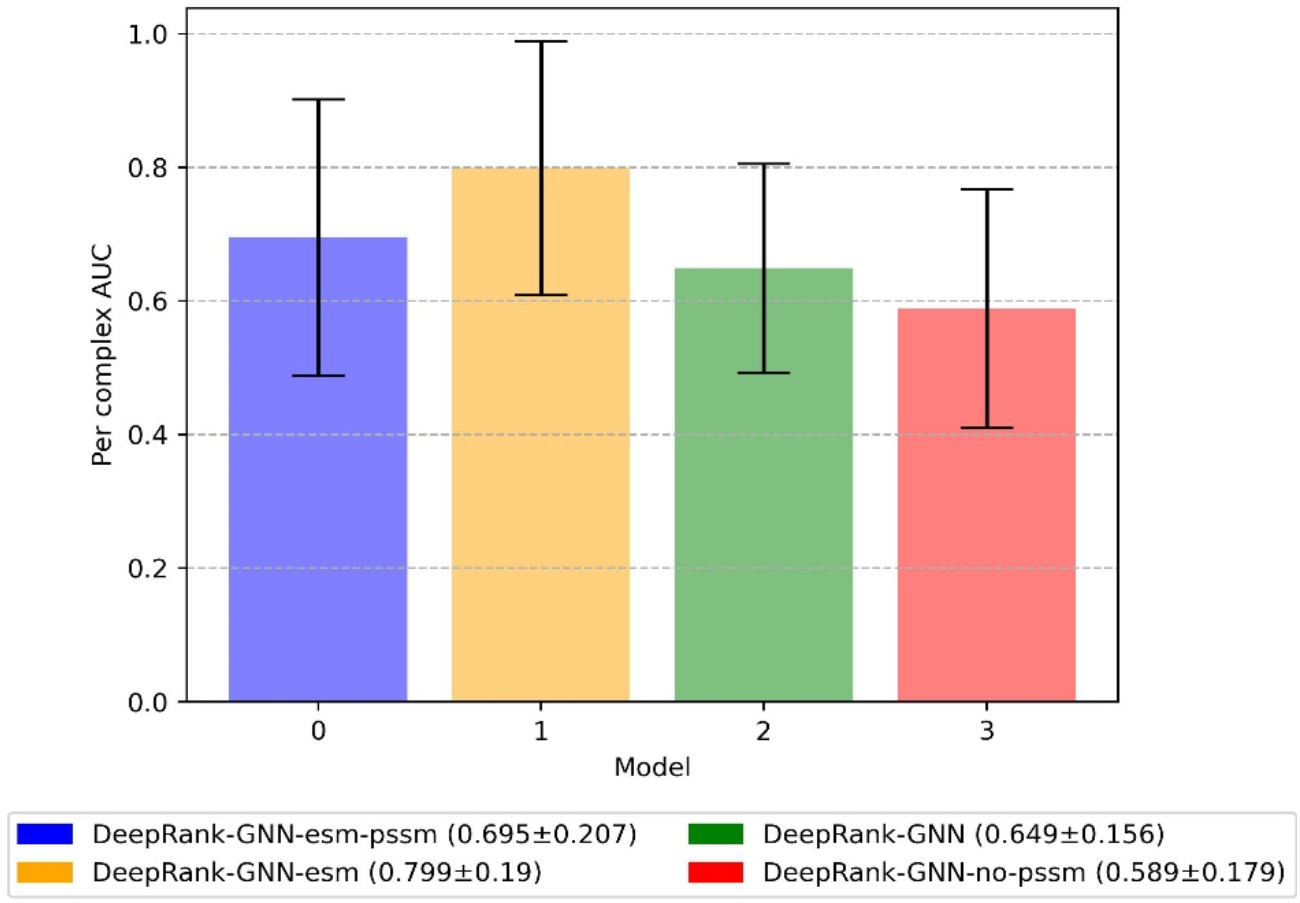
Average per complex AUC value with standard deviation for the four models on the BM5 test set.

In the CAPRI score set, the DeepRank-GNN-esm model exhibits the highest predictive performance (69.2% top50), in par with iScore and GNN-Dove, even surpassing the DeepRank-GNN-esm-pssm model. Within the top 50 ranks, the DeepRank-GNN-esm model successfully identifies correct complex conformations for nine out of 13 targets. DeepRank-GNN-esm-pssm, DeepRank-GNN, DeepRank and HADDOCK closely follow with success rates of 61.5% (Table 3). Notably, six graph-based approaches, including DeepRank-GNN and its three derivatives, iScore, and GNN_Dove, consistently demonstrate higher success rates at earlier ranks, distinguishing themselves from the other methods. iScore performs well at rank 1 and rank 3, while the DeepRank-GNN-esm model excels at later ranks. Combining the DeepRank-GNN-esm model with iScore predictions could potentially yield even better predictions.

#### 3.2.3 Application of DeepRank-GNN-esm model for discriminating physiological from non-physiological interfaces

Next to scoring, we also trained our models on the task of discriminating biological interfaces from crystallographic ones using the MANY dataset for training and the DC dataset as independent test set. During the training process, we monitored the loss curves and calculated AUC values (Supplementary Figure 6). In contrast to the scoring data set, in this case both the training and test datasets are perfectly balanced and the accuracy is therefore a suitable metric for comparing the performance of the models. Both DeepRank-GNN-esm-pssm and DeepRank-GNN-esm achieve an accuracy of 0.83 (Table 4), comparable to the reported accuracy of DeepRank-GNN (0.82) on the DC test dataset. Our results demonstrate the effectiveness of ESM features in accurately identifying true biological interfaces from crystal artifacts.

**Table 4.** Comparison of model performances on the CAPRI Score set

| Model<sup>a</sup> | Accuracy (%) |
| --- | --- |
| PISA | 79 |
| PRODIGY-Crystal | 74 |
| DeepRank | 86 |
| Model1 | 83 |
| Model2 | 83 |
| Model3 | 82 |

a) Model 1: DeepRank-GNN-esm-pssm, Model 2: DeepRank-GNN-esm, Model 3: DeepRank-GNN.

### 3.3 Computational speed

The gain in efficiency of using ESM-2 embeddings compared to traditional PSSM profiles is significant: Generating a single PSSM profile requires a non-redundant protein database of 176GB and takes approximately two hours when computed on a single core. In contrast, generating ESM-2 embeddings for the same sequence only requires a pre-trained model of 2.5GB size and takes approximately 5 seconds. This represents a more than 100-fold increase in efficiency compared to the PSSM generation process. These findings are supported by data presented in Supplementary Table S2, which highlights the efficiency gains achieved by ESM-2 embeddings. Our data also indicate that there is no significant increase in model inference time associated with the use of protein language model features.

## 4. Conclusions

By integrating language model-based features into our existing deep learning framework, we have developed the DeepRank-GNN-esm algorithm, which enhances the scoring of protein-protein complexes. Adding the language model embeddings results in an increased performance. Overall, our findings on two PPI-related tasks (scoring and discrimination of biological interfaces) suggest that PSSM features can be replaced by EMS-2 embeddings, leading to improved model performance. This comes with the advantage of bypassing the requirement of pre-generating the PSSM, a cumbersome and computationally more expensive process, while maintaining or even slightly improving performance. This opens new avenues for future research, particularly in the case of antibody-antigen complexes or protein-peptide complexes where PSSM profiles may not be applicable. To the best of our knowledge, DeepRank-GNN-esm is the first method to apply protein language models in graph neural networks for protein-protein models evaluation tasks.

## 5. Availability

Software and related data are freely available on GitHub at https://github.com/DeepRank/DeepRank-GNN-esm BM5 docking models and CAPRI score set, MANY and DC dataset used in training and evaluation can be downloaded from https://data.sbgrid.org/dataset/843/.

## Supplementary Material

### Supplementary Method 1 Details of ESM-2 embeddings calculations

We first extracted the sequence for each chain (A, B) in all protein-protein complexes. To compute esm-2 embeddings for those sequences in bulk, we used the python script (extract.py) provided at https://github.com/facebookresearch/esm with the command below:

~~~
python extract.py esm2_t33_650M_UR50D all.fasta ../ -repr_layers 0 32 33 --include mean per_tok
~~~

In the above command, “`esm2_t33_650M_UR50D`” specifies the pre-trained model to be used for the embeddings. We chose to include embeddings from layers 0, 32, and 33 by using the ‘ `reprlayers`’ option. The ‘ `include`’ option was used to generate an averaged embedding per amino acid per layer over the entire protein sequence. This means that for each layer, the script computed an embedding for every amino acid in the sequence and then averaged those embeddings to obtain a single representation per amino acid per layer. As a result of this process, the script generated one. `pt` file per FASTA sequence. Each. `pt` file contains the embeddings for each residue in the protein sequence. Subsequently, these generated embeddings were associated with nodes in protein interface graphs and incorporated into the HDF5 files used by DeepRank-GNN.

### Supplementary Method 2 Definition of the six machine learning metrics used for assessing the performance

1. Precision measures the accuracy of positive predictions made by the model as such:

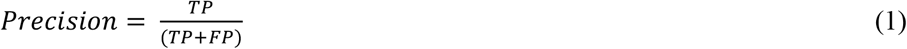
2. MCC (Matthews’s correlation coefficient) quantifies the overall quality of binary classifications, considering true and false positives and negatives as such:

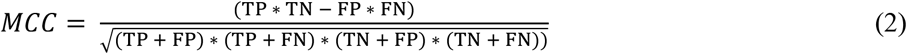
3. Recall calculates the proportion of actual positives correctly identified by the model as such:

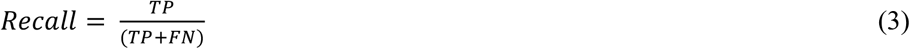
4. F1 Score combines precision and recall to provide a balanced measure of model perfo rmance as such:

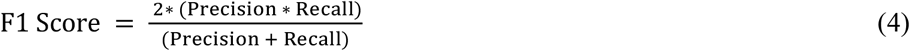
5. R^2^ score measures the proportion of the variance in the dependent variable that can be explained by the independent variable(s) as such:

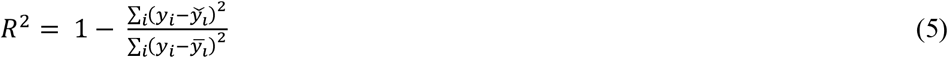
6. The Pearson correlation coefficient quantifies the strength and direction of t he linear relationship between two variables as such

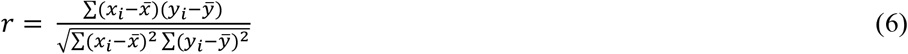

**Supplementary Table S1.** Data composition of the training, evaluation, and test sets

| **Dataset** | **Number of complexes** | **Total** | **Positive** | **Negative** | **Percentage of positive models** |
| --- | --- | --- | --- | --- | --- |
| BM5_train | 127 | 2298799 | 166893 | 2131906 | 0.0726 |
| BM5_eval | 127 | 559664 | 40373 | 519291 | 0.0721 |
| BM5_test | 15 | 338995 | 19459 | 319536 | 0.0574 |
| CAPRI_test | 13 | 16593 | 1968 | 14625 | 0.1186 |

**Supplementary Table S2.** Computing time requirements comparison between PSSM and esm features

| **Model** | **Feature Generation Time per model** | **Average Training Time per epoch (hour)** | **Average Inference Time per epoch (hour)** | **Average Inference Time per model (second)** |
| --- | --- | --- | --- | --- |
| DeepRank-GNN-esm-pssm | 2 hours | 30.97 | 2.52 | 0.016 |
| DeepRank-GNN-esm | 5 seconds | 29.30 | 2.45 | 0.015 |
| DeepRank-GNN | 2 hours | 24.80 | 2.35 | 0.015 |
| DeepRank-GNN-no-pssm | 5 seconds | 8.64 | 1.52 | 0.009 |

\*All models were trained on a GeForce GTX 1080 Ti.

**Supplementary Figure S1.**
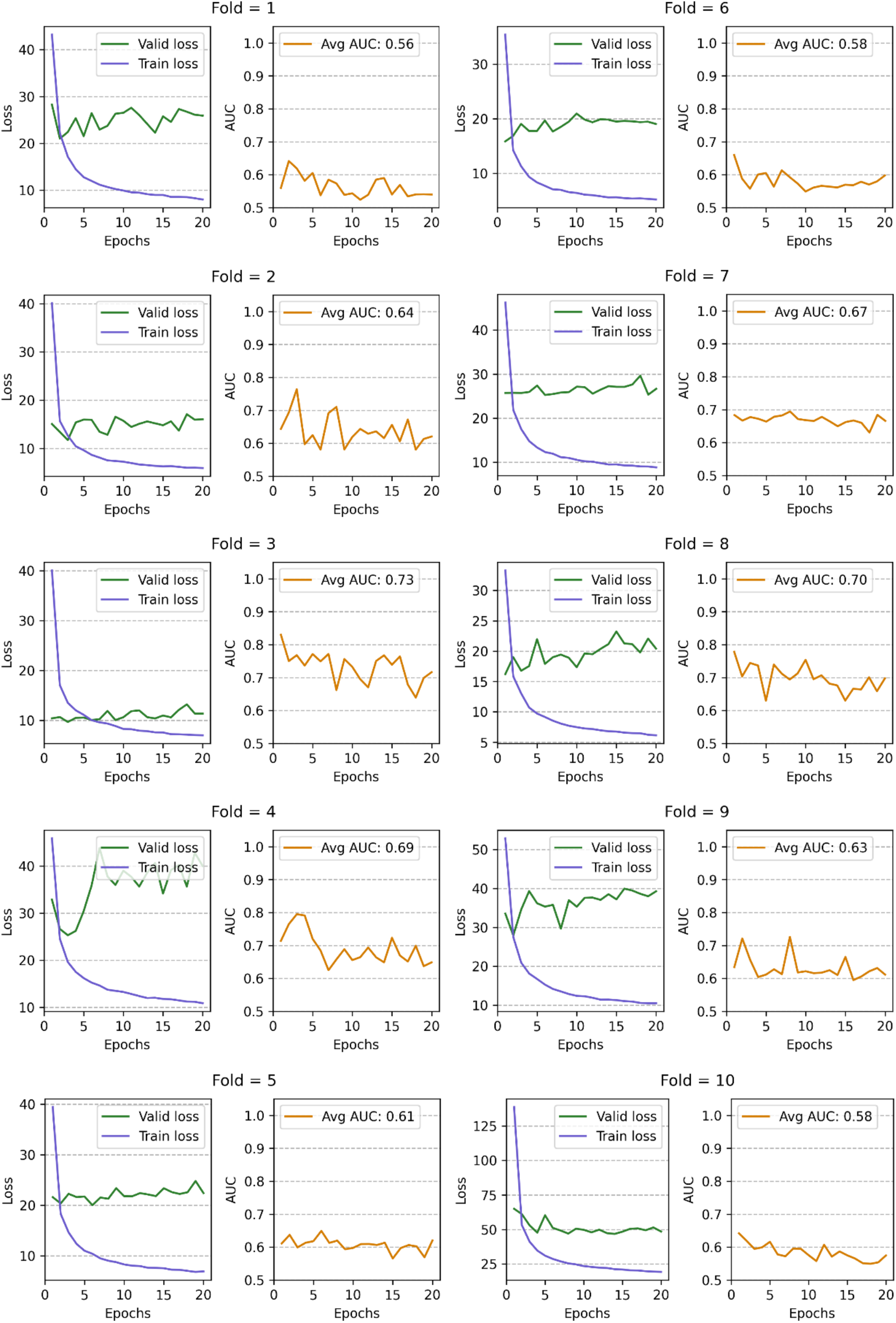
Losses and AUC curves for the DeepRank-GNN-esm-pssm models during cross-validation

**Supplementary Figure S2.**
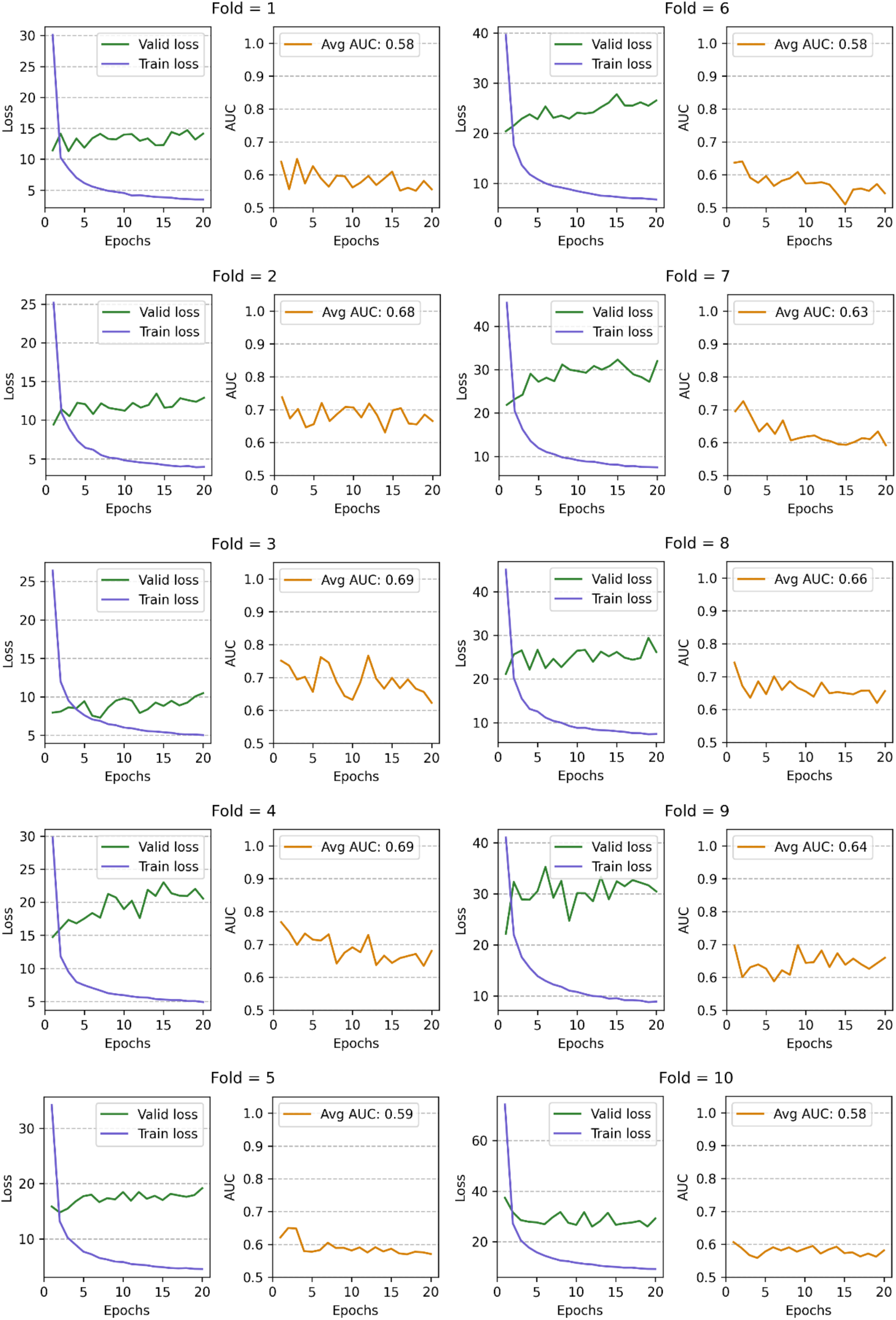
Losses and AUC curves for the DeepRank-GNN-esm model during cross-validation

**Supplementary Figure S3.**
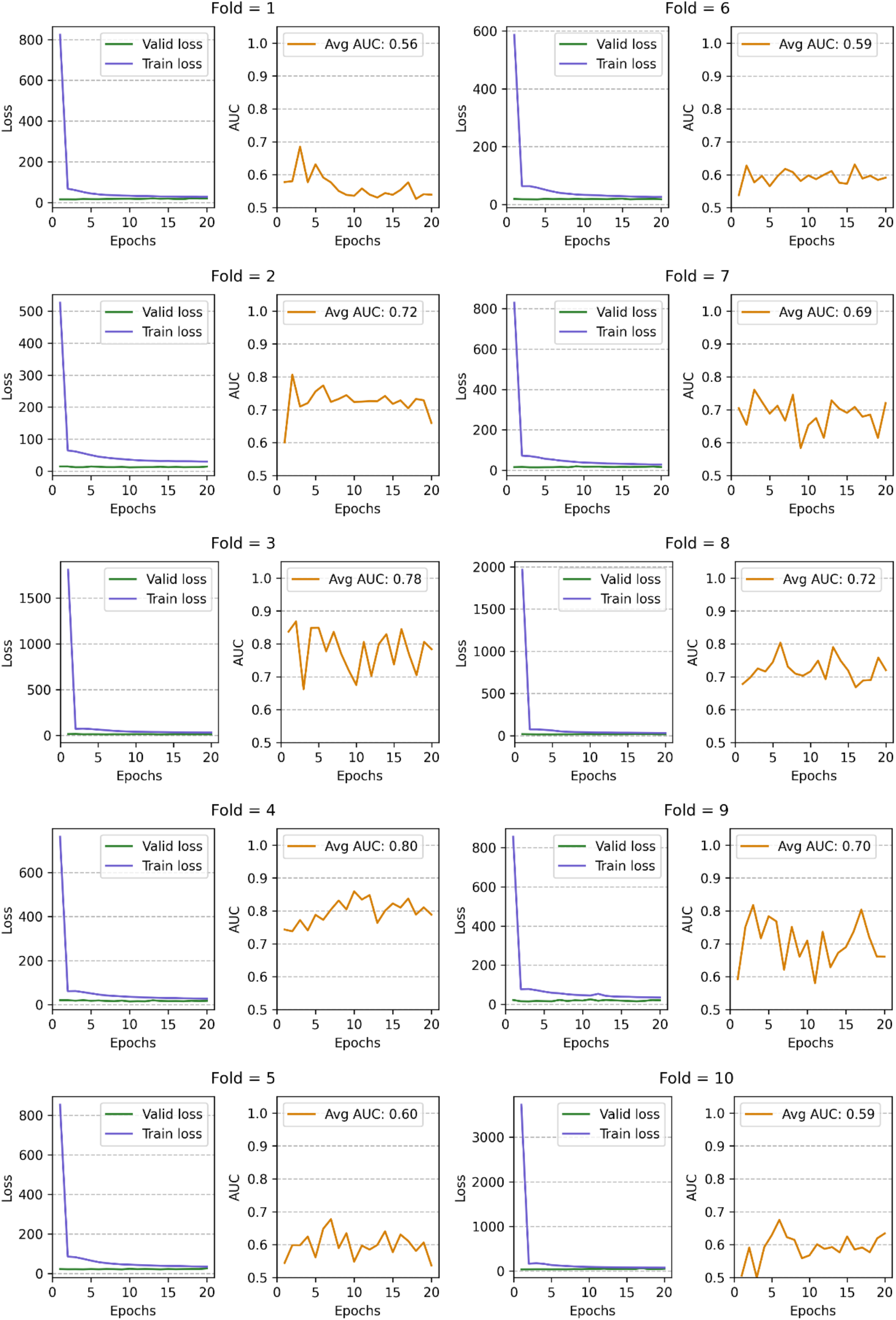
Losses and AUC curves for the DeepRank-GNN-no-pssm model during cross-validation

**Supplementary Figure S4.**
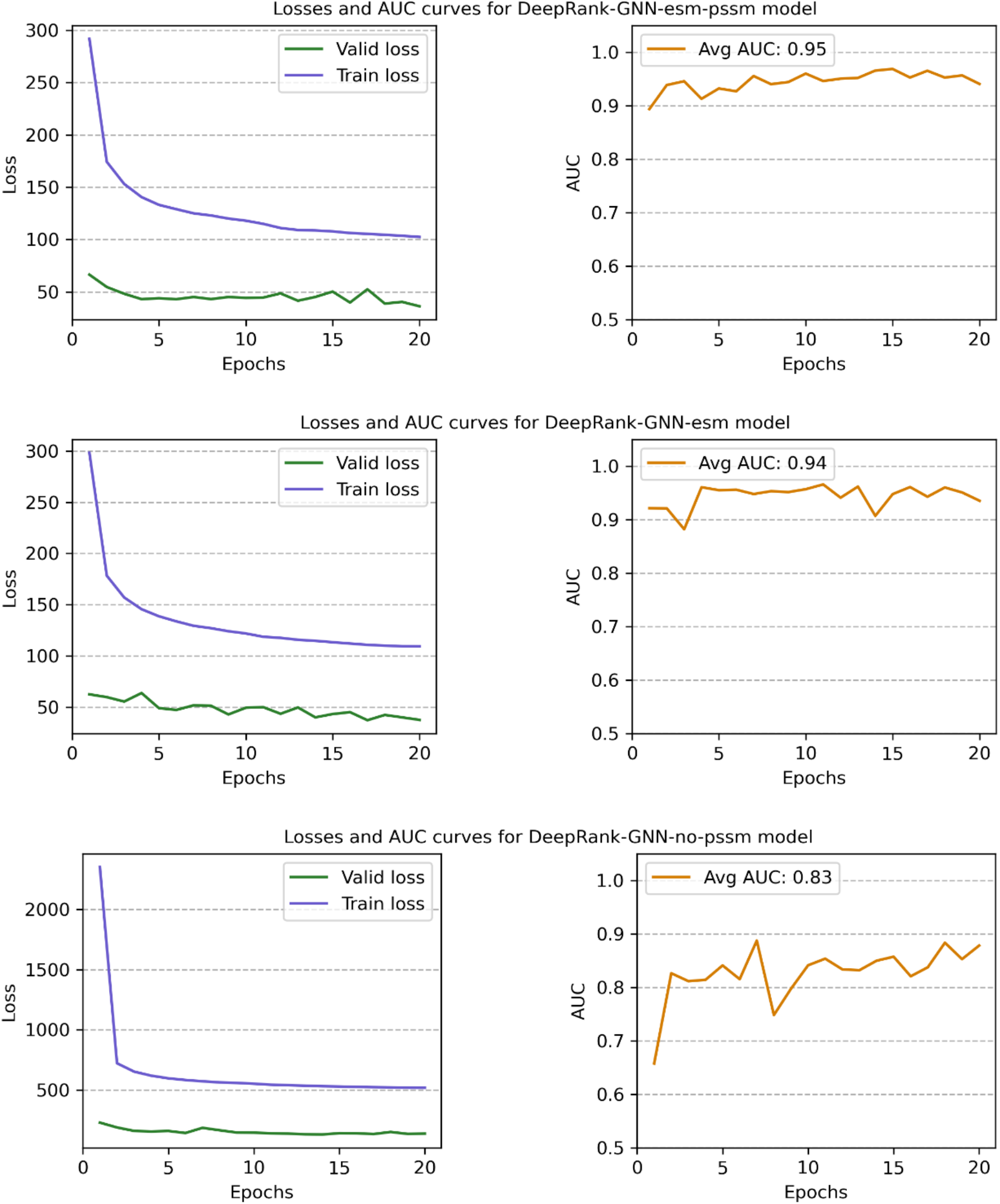
Losses and AUC curves for three final models

**Supplementary Figure S5.**
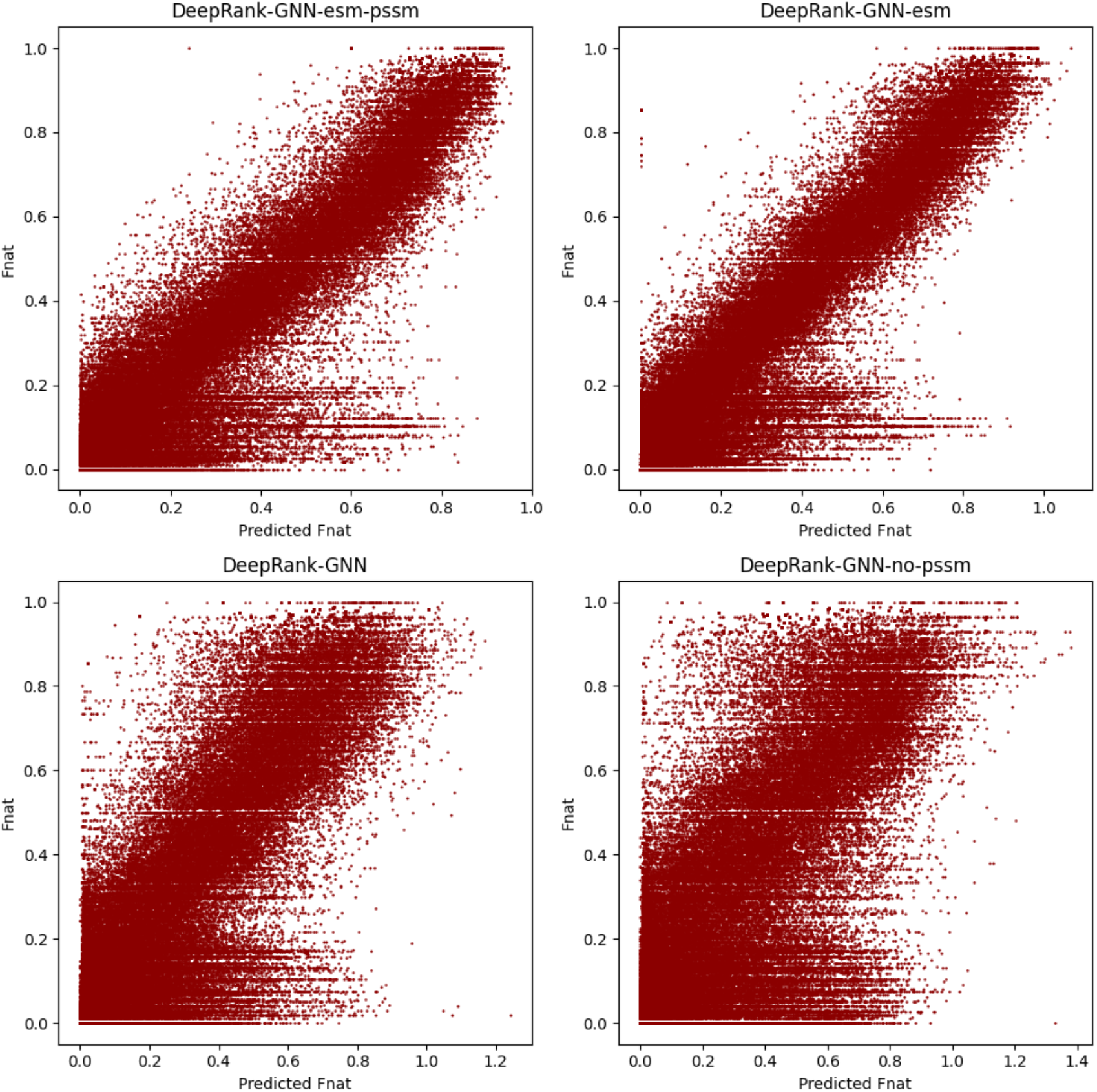
Scatter plots of Fnat versus predicted Fnat for the four models on BM5 evaluation set

**Supplementary Figure S6.**
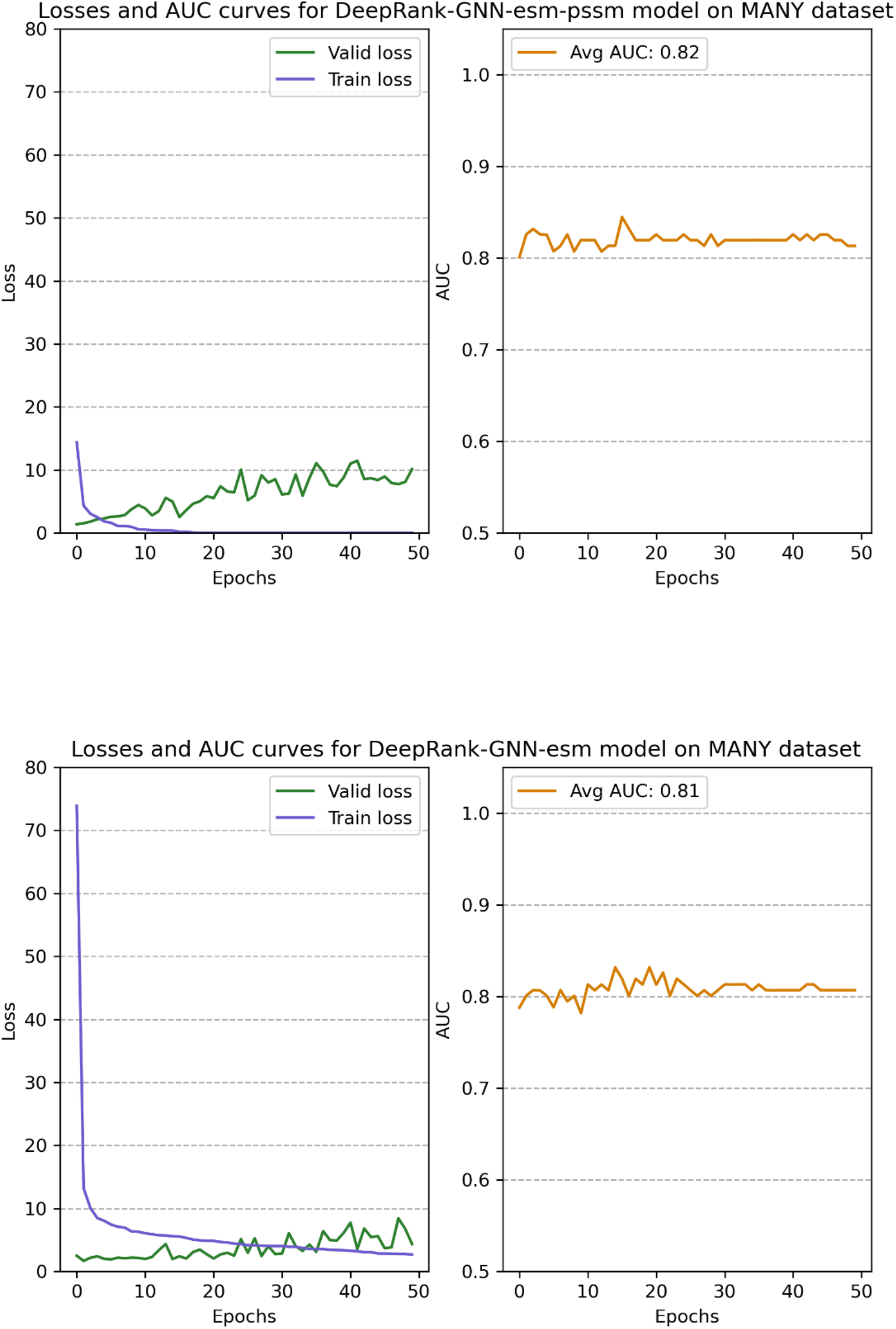
Losses and AUC curves of two models trained on MANY dataset

